# A revised biosynthetic pathway for the cofactor F_420_ in bacteria

**DOI:** 10.1101/470336

**Authors:** Ghader Bashiri, James Antoney, Ehab N. M. Jirgis, Mihir V. Shah, Blair Ney, Janine Copp, Stephanie M. Stutely, Sreevalsan Sreebhavan, Brian Palmer, Martin Middleditch, Nobuhiko Tokuriki, Chris Greening, Edward N. Baker, Colin Scott, Colin J. Jackson

**Author notes:** These authors contributed equally to this work (Ghader Bashiri, James Antoney). Corresponding author contact details: Ghader Bashiri, Colin Jackson, Colin Scott.

## Abstract

Cofactor F_420_ plays critical roles in primary and secondary metabolism in a range of bacteria and archaea as a low-potential hydride transfer agent. It mediates a variety of important redox transformations involved in bacterial persistence, antibiotic biosynthesis, pro-drug activation and methanogenesis. However, the biosynthetic pathway for F_420_ has not been fully elucidated: neither the enzyme that generates the putative intermediate 2-phospho-*L*-lactate, nor the function of the FMN-binding C-terminal domain of the γ-glutamyl ligase (FbiB) in bacteria are known. Here we show that the guanylyltransferases FbiD and CofC accept phosphoenolpyruvate, rather than 2-phospho-*L*-lactate, as their substrate, leading to the formation of the previously uncharacterized intermediate, dehydro-F_420_-0. The C-terminal domain of FbiB then utilizes FMNH2 to reduce dehydro-F_420_-0, which produces mature F_420_ species when combined with the γ-glutamyl ligase activity of the N-terminal domain. This new insight has allowed the heterologous expression F_420_ from a recombinant F_420_ biosynthetic pathway in *Escherichia coli*.

---

Cofactor F_420_ is a deazaflavin that acts as a hydride carrier in diverse redox reactions in both bacteria and archaea^1,2^. While F_420_ structurally resembles the flavins FMN and FAD, it functions as an obligate two-electron hydride carrier and hence is functionally analogous to the nicotinamides NAD^+^ and NADP^+3^. The lower reduction potential of the F_420_, relative to the flavins, results from the substitution of N5 of the isoalloxazine ring in the flavins for a carbon in F_420_^4,5^. Originally characterized from methanogenic archaea in 1972^4,5^, F_420_ is an important catabolic cofactor in methanogens and mediates key one-carbon transformations of methanogenesis^6^. F_420_ has since been shown to be synthesized in a range of archaea and bacteria^1,2,7,8^. In *Mycobacterium tuberculosis*, the causative agent of tuberculosis, F_420_ has been shown to contribute to persistence^9,10^ and to activate the new clinical antitubercular prodrugs delamanid and pretomanid^11^. There are also growing numbers of natural products that have been shown to be synthesized through F_420_-dependent pathways, including tetracyclines^12^, lincosamides^13^, and thiopeptides^14^. F_420_-dependent enzymes have also been explored for bioremediation and biocatalytic applications^15,16^.

The currently accepted F_420_ biosynthetic pathway consists of two branches^2^. In the first branch, tyrosine is condensed with 5-amino-6-ribitylamino-2,4[1*H*,3*H*]-pyrimidinedione from the flavin biosynthetic pathway to generate the deazaflavin chromophore Fo (7,8-dide-methyl-8-hydroxy-5-deazariboflavin) *via* the activity of the two-domain Fo synthase FbiC, or the CofG/H pair (where ‘Fbi’ refers to mycobacterial proteins and ‘Cof’ refers to archaeal homologs). In the second branch, it has been hypothesized that a 2-phospho-*L*-lactate guanylyltransferase (CofC in archaea and the putative enzyme FbiD in bacteria) catalyzes the guanylylation of 2-phospho-*L*-lactate (2-PL) using guanosine-5’-triphosphate (GTP), yielding *L*-lactyl-2-diphospho-5’-guanosine (LPPG)^17^. The two branches then merge at the reaction catalyzed by the transferase FbiA/CofD, where the 2-phospho-*L*-lactyl moiety of LPPG is transferred to Fo, forming F_420_-0^18,19^. Finally, the γ-glutamyl ligase (FbiB/CofE) catalyzes the poly-glutamylation of F_420_ to generate mature F_420_, with poly-γ-glutamate tail lengths of ∼2-8, depending on species^20,21^.

There are three aspects of the F_420_ biosynthetic pathway that require clarification. First, the metabolic origin of 2-PL, the proposed substrate for CofC, is unclear. It has been assumed that a hypothetical kinase (designated CofB) phosphorylates *L*-lactate to produce 2-PL^22^. However, no such enzyme for the production of 2-PL has been identified in bacteria or archaea, and our genomic analysis of F_420_ biosynthesis operons has failed to identify any candidate enzymes with putative *L*-lactate kinase activity^2^. Second, the existence of FbiD has only been inferred through bioinformatics and genetic knockout studies and the enzyme has not been formally characterized^23,24^. Finally, the bacterial γ-glutamyl ligase FbiB is a two-domain protein^20^, in which the N-terminal domain is homologous to other F_420_-γ-glutamyl ligases (including the archaeal equivalent, CofE) and the C-terminal domain adopts an FMN-binding nitroreductase (NTR) fold^20^. Although both domains are required for full γ-glutamyl ligase activity, no function has been associated with either the C-terminal domain or the FMN cofactor, given no redox reactions are known to be involved in F_420_ biosynthesis.

Here we demonstrate that 2-PL is not required for F_420_ biosynthesis in bacteria and instead phosphoenolpyruvate (PEP), an abundant intermediate of glycolysis and gluconeogenesis, is incorporated into F_420_. We show that PEP guanylylation is catalyzed by the CofC and FbiD enzymes that were previously thought to act upon 2-PL. In bacteria, the incorporation of PEP in the pathway results in the production of the previously undetected intermediate dehydro-F_420_-0, which is then reduced by the C-terminal domain of FbiB alongside poly-glutamylation. These findings result in a substantially revised pathway for F_420_ biosynthesis and have allowed us to heterologously produce functional F_420_ in engineered *Escherichia coli*, an organism that does not normally produce F_420_, at levels comparable to some native F_420_-producing organisms.

## Results

### FbiD/CofC use phosphoenolpyruvate, rather than 2-phospho-*L*-lactate, as a substrate

The archaeal enzyme CofC has previously been suggested to catalyze the guanylylation of 2-PL to produce LPPG during F_420_ biosynthesis (Fig. 1a)^25^. Another study, using transposon mutagenesis, has shown that MSMEG_2392 of *Mycobacterium smegmatis* is essential in the biosynthesis of F_420_ from Fo^23^. We have recently shown that homologs of this gene have sequence homology to CofC and belong to operons with other validated F_420_ biosynthetic genes in a wide variety of bacteria^2^. In keeping with the bacterial naming system, we refer to this enzyme as FbiD. To test the function of this putative bacterial FbiD, we cloned the Rv2983 gene from *M. tuberculosis*^26^ into a mycobacterial expression vector and purified heterologously expressed *Mtb*-FbiD from *M. smegmatis* mc^2^4517 host cells. We also expressed and purified *Mtb*-FbiA (the enzyme thought to transfer the 2-phospho-*L*-lactyl moiety of LPPG to Fo to produce F_420_-0)^18^ to use in coupled HPLC-MS enzymatic assays with *Mtb*-FbiD. Surprisingly, we found that when *Mtb*-FbiD and *Mtb*-FbiA were included in an assay with 2-PL, GTP (or ATP) and Fo, no product was formed (Fig. 1b). We then tested whether *Mtb*-FbiA and CofC from *Methanocaldococcus jannaschii* (*Mj*-CofC) could catalyze F_420_-0 formation under the same conditions, which again yielded no product (Fig. 1b).

**Fig. 1.**
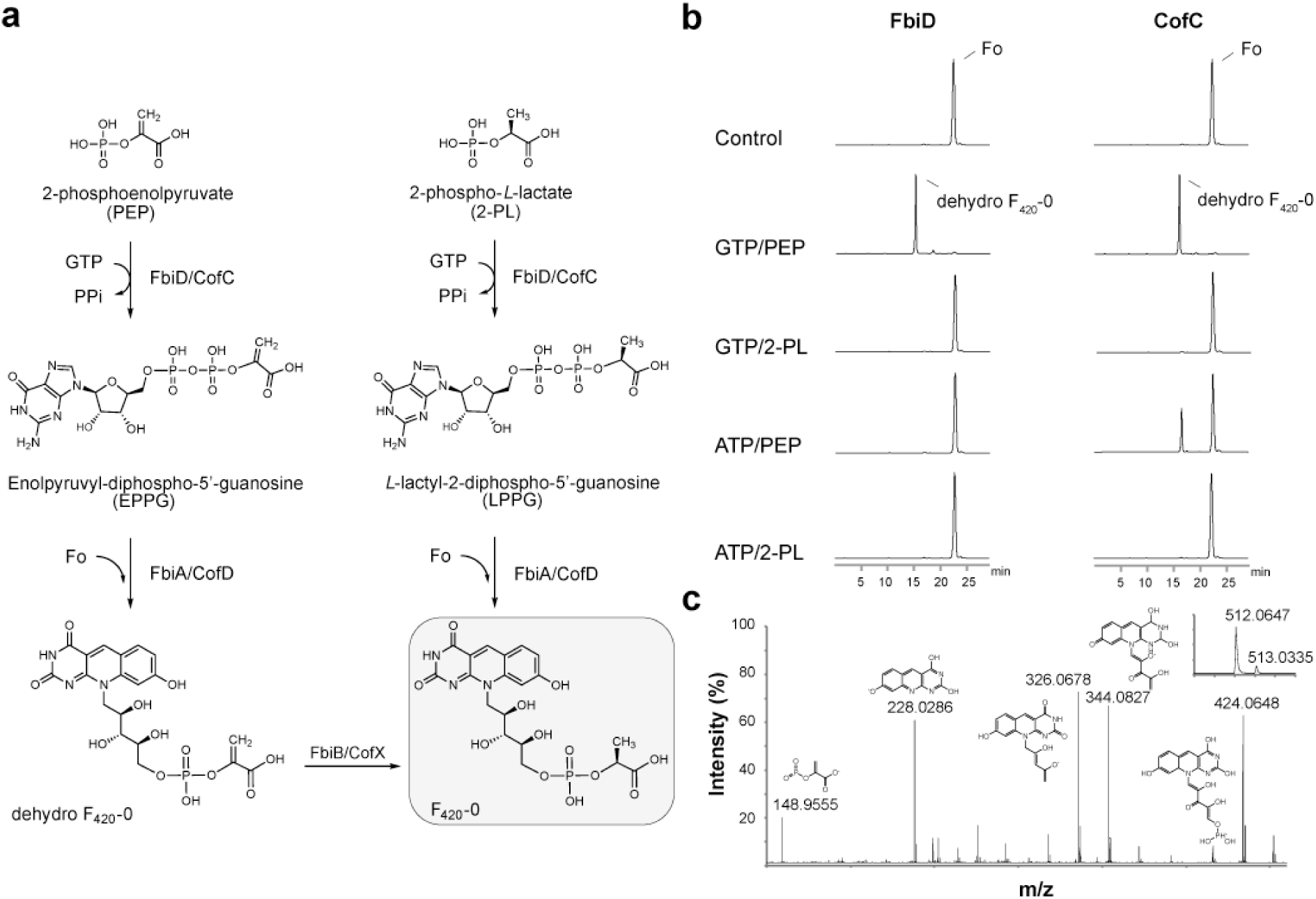
Phosphoenolpyruvate (PEP) is an intermediate in the formation of dehydro-F_420_-0. **(a)** Production of F_420_-0 in our revised biosynthesis pathway (left) compared to the cur-rently accepted pathway (right). **(b)** Coupled-reaction HPLC assays showing that both *Mtb*-FbiD and *Mj*-CofC enzymes use PEP to produce dehydro-F_420_-0. **(c)** Tandem mass spectral identification of dehydro-F_420_-0. MS/MS fragmentation of dehydro-F_420_-0, showing fragment ions with their corresponding structures. The inset displays the observed spectrum of the parent molecule (expected monoisotopic *m*/*z* 512.0711 [M-H]^-^).

Although 2-PL is hypothesized to be an intermediate in F_420_ biosynthesis, this has never been experimentally confirmed in bacteria. Additionally, no enzyme capable of phosphorylating *L*-lactate to 2-PL has been identified in F_420_ producing organisms, despite considerable investigation^22^. 2-PL has been little studied as a metabolite and is only known to occur as a by-product of pyruvate kinase activity^27^. 2-PL has not been implicated as a substrate in any metabolic pathway outside the proposed role in F_420_ biosynthesis; rather it has been shown *in vitro* to inhibit several enzymes involved in glycolysis and amino acid biosynthesis^28-30^. Our inability to detect activity with 2-PL led us to consider alternative metabolites that could potentially substitute for 2-PL, namely the structurally analogous and comparatively abundant molecule phosphoenolpyruvate (PEP)^31^ (Fig. 1a).

While there was no activity when 2-PL was used in the FbiD/CofC:FbiA coupled assays, when these enzymes were incubated with PEP, GTP (or ATP) and Fo, a previously unreported intermediate in the F_420_ biosynthesis pathway, which we term ‘dehydro-F_420_-0’, was produced (Fig. 1b). The identity of this compound, which is identical to F_420_-0 except for a methylene group in place of the terminal methyl group, was verified by tandem mass spectrometry (Fig. 1c). The only difference that we observed between *Mtb*-FbiD or *Mj*-CofC was that while *Mtb*-FbiD exclusively utilizes GTP to produce dehydro-F_420_-0, *Mj*-CofC can also catalyze the reaction with ATP, albeit to a lesser extent (Fig. 1b). Interestingly, in our experiments the FbiD/CofC enzymes were only active in the presence of FbiA. This was not unexpected given that the inferred intermediate (enolpyruvyl-diphospho-5’-guanosine; EGGP) is expected to be unstable^22^ (Fig. 1a).

To understand the molecular basis of PEP recognition by *Mtb*-FbiD, we crystallized the protein and solved the structure by selenium single-wavelength anomalous diffraction (Se-SAD), and then used this selenomethionine-substituted structure to obtain the native FbiD structure by molecular replacement. The latter was then refined at 1.99 Å resolution (R/Rfree=0.19/0.22) (SI Table 2). As expected, *Mtb*-FbiD adopts the same MobA-like nucleoside triphosphate (NTP) transferase family protein fold as CofC: a central 7-stranded β-sheet (six parallel strands and one antiparallel), with α-helices packed on either side (Fig. 2a). Superposition of CofC from *Methanosarcina mazei* (PDB code 2I5E) on to *Mtb*-FbiD gives a root mean square difference (rmsd) of 1.85 Å over 181 Cα atoms, with 25.4% sequence identity, establishing clear structural homology.

**Fig. 2.**
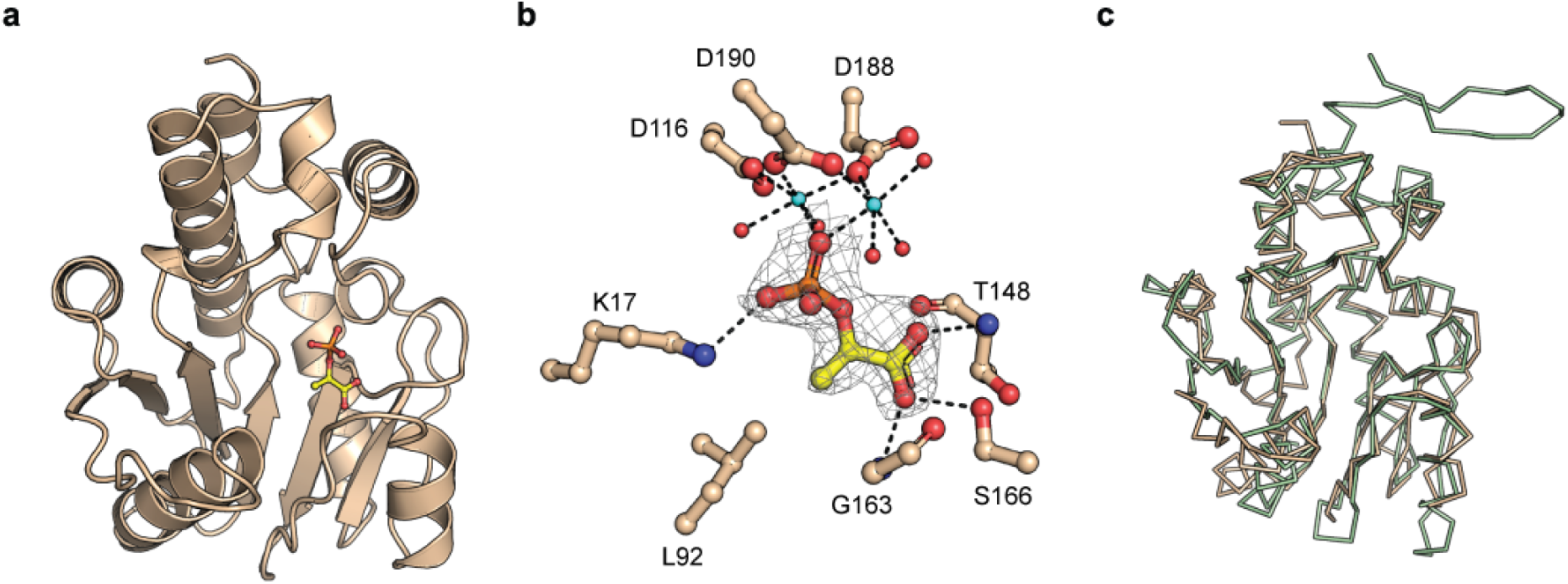
Crystal structure of *Mtb*-FbiD. **(a)** Cartoon representation of the *Mtb*-FbiD structure in complex with PEP, shown as a ball-and-stick model. **(b)** The phosphate group of PEP binds to three aspartic acid side chains through two Mg^2+^ ions (shown in cyan). PEP is shown in 2*F*_o_ - *F*_c_ omit density contoured at 2.0 σ, and drawn as ball-and-stick model. Water molecules are shown as red spheres and hydrogen bond interactions are outlined as dashed lines. **(c)** Superposition of *Mtb*-FbiD (wheat ribbon) and *Mj*-CofC (green ribbon), indicating 1.85 Å rmsd over 181 superimposed Cα. The protruding hairpin in the *Mj*-CofC structure is involved in dimer formation, which is absent in *Mtb*-FbiD.

We also soaked PEP into pre-formed FbiD crystals to obtain an FbiD-PEP complex (2.18 Å resolution, R/R_free_=0.22/0.26). FbiD has a cone-shaped binding cleft with a groove running across the base of the cone, formed by the C-terminal end of the central β-sheet (Fig. 2a). PEP binds in the cleft with its phosphate group anchored through two Mg^2+^ ions to three acidic side chains (D116, D188 and D190) (Fig. 2b). The PEP carboxylate group is hydrogen bonded to the hydroxyl group of S166 and the main chain NH groups of T148 and G163. All PEP binding residues are conserved in the CofC protein of *M. mazei* (PDB code 2I5E) (Fig. 2c and Fig. S2), consistent with the enzymatic assays that showed PEP is the substrate for archaeal CofC, as well as FbiD. In the GTP-bound structure of *E. coli* MobA^32^ (PDB code 1FRW), GTP binds in a characteristic surface groove, providing a structural framework for substrate binding and catalysis. In our enzyme assays, neither FbiD nor CofC showed activity in the absence of FbiA. Moreover, we did not observe GTP binding in either our co-crystallization or differential scanning fluorimetry experiments. We speculate that the GTP binding site is not formed until FbiD/CofC interacts with FbiA/CofD, enabling catalysis to proceed through to formation of dehydro-F_420_-0. This may provide an advantage by producing EPPG/EPPA only when both proteins are available, thereby overcoming the issue of intermediate instability.

### The C-terminal domain of FbiB catalyzes reduction of dehydro-F_420_-0 to F_420_-0

Dehydro-F_420_-0 would yield F_420_-0 upon reduction of the terminal methylene double bond. However, no masses corresponding to F_420_-0 were identified in any of the LC-MS traces from the FbiD:FbiA coupled assays, suggesting that an enzyme other than FbiD or FbiA catalyzes dehydro-F_420_-0 reduction. We have previously shown that full-length FbiB consists of two domains: an N-terminal domain that is homologous to the archaeal γ-glutamyl ligase CofE^21,33^, and a C-terminal domain of the NTR fold^34^ that binds to FMN and has no known function^20^, but is essential for extending the poly-γ-glutamate tail.

We tested whether FbiB could use dehydro-F_420_-0 as a substrate with a three enzyme assay in which FbiB and *L*-glutamate were added to the FbiD:FbiA coupled assay. *Mtb-*FbiB was observed to catalyze the addition of *L*-glutamate residues to dehydro-F_420_-0, forming de-hydro-F_420_ species with one ([M+H]^+^, *m*/*z* of 643.40) and two ([M+H]^+^, *m*/*z* of 772.40) glutamate residues (Fig. S1). We then tested the hypothesis that the orphan function of the FMN-binding C-terminal domain could in fact be a dehydro-F_420_-0 reductase. We used a four-enzyme *in vitro* assay where *E. coli* NAD(P)H:flavin oxidoreductase^35^ (Fre), FMN, NADH and 10 mM dithiothreitol (to maintain reducing conditions and generate reduced FMNH2) were added to the FbiD:FbiA:FbiB assay and the reaction was performed in anaerobic conditions (to prevent reaction of FMNH2 with oxygen). We found that F_420_-1, *i.e.* the fully reduced and mature glutamylated cofactor, was produced; but only in the presence of both FbiB and Fre/FMNH2 (Fig. 3a). Thus, dehydro-F_420_-0 is a *bona fide* metabolic intermediate that can be converted to mature F_420_ by FbiB in an FMNH_2_-dependent fashion. These results demonstrate that bacterial FbiB is a bifunctional enzyme, functioning as a dehydro-F_420_-0 reductase and as a γ-glutamyl ligase (Fig. 3c).

**Fig. 3.**
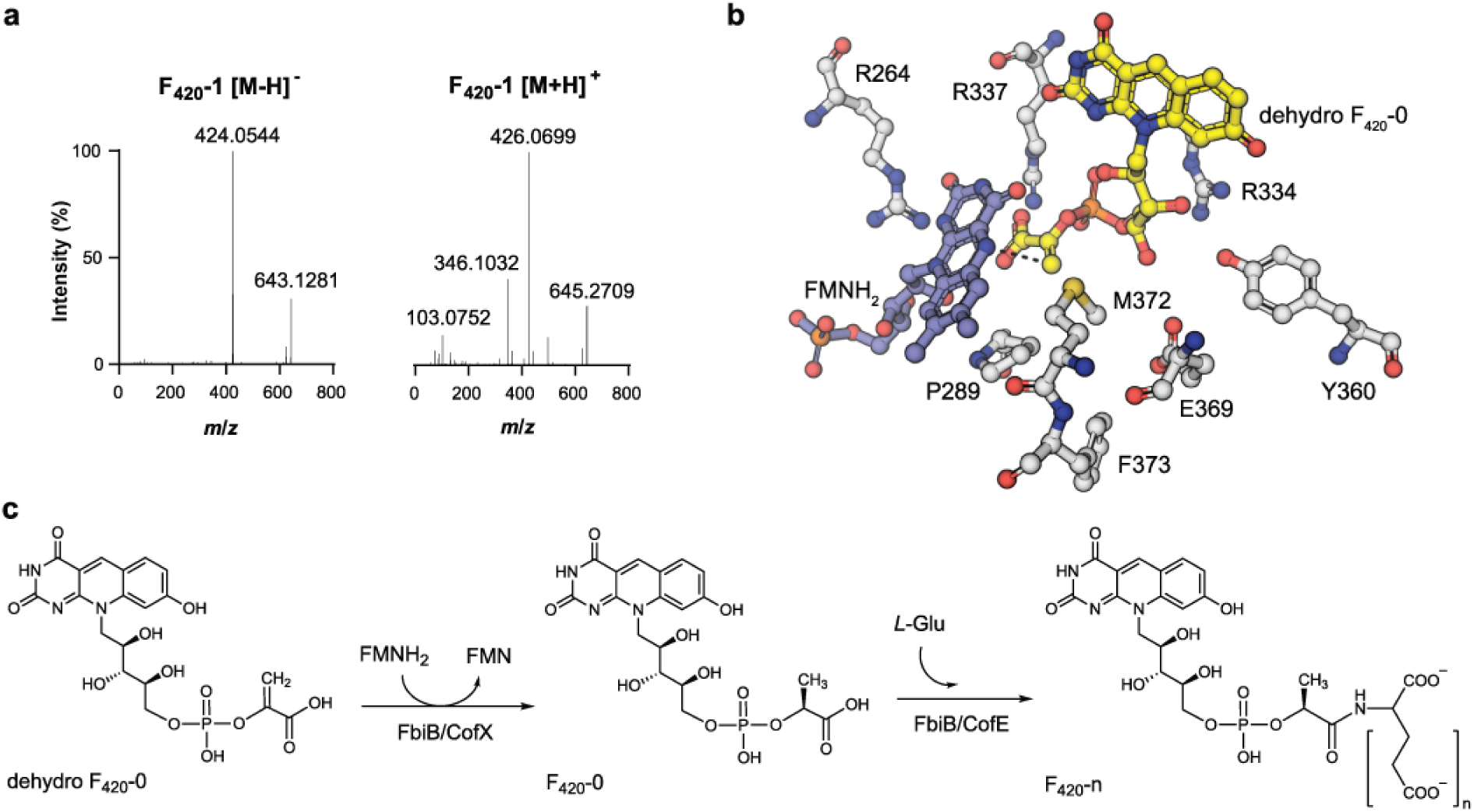
*Mtb*-FbiB catalyzes reduction of dehydro-F_420_-0. **(a)** F_420_-1 is produced in FbiD:FbiA:FbiB coupled assays in the presence of Fre/FMNH2 and *L*-glutamate. MS/MS confirmation of F_420_-1 in both negative (643.12811, [M-H]^-^) and positive (645.27094, [M+H]^+^) modes. **(b)** Docking of FMNH2 and dehydro-F_420_-0 into the crystal structure of FbiB C-terminal domain. The methylene group of the enolpyruvyl moiety sits in a pocket made up of M372 and P289 while the carboxylate hydrogen bonds with R337. The methylene double bond sits planar above the isoalloxazine ring of FMNH2 at an appropriate distance (3.6 Å, shown by dashed line) and oriented for a hydride transfer to the *Si* face of the methylene bond, accounting for the observed (*S*)-lactyl moiety of F_420_. **(c)** *Mtb*-FbiB is a bifunctional enzyme catalyzing the reduction of dehydro-F_420_-0 and its poly-glutamylation to form F_420_-n.

When the previously published crystal structures of *Mtb*-FbiB are analyzed in the context of these results, the molecular basis for this activity becomes clear. Crystal structures with both F_420_ and FMN bound have been separately solved and when these are overlaid, it is apparent that the FMN molecule is ideally situated to transfer a hydride to the terminal methylene of dehydro-F_420_-0 (assuming deydro-F_420_-0 binds in a similar fashion to F_420_). Interestingly, the phospholactyl group of F_420_ appears to be disordered in these crystal structures, suggesting it may adopt multiple conformations. To test this, we docked and minimized dehydro-F_420_-0 and FMNH2 simultaneously using the OPSL3a forcefield. The results show that in this ternary complex the two molecules can adopt ideal positions and orientations for the reduction of dehydro-F_420_-0 (Fig. 3b). The methylene group of dehydro-F_420_-0 is accommodated by a small hydrophobic pocket mostly comprised of P289 and M372 allowing it to be positioned above the N5 atom of FMNH2, in a plausible Michaelis complex for hydride transfer. We therefore suggest that the phosphoenolpyruvyl (analogous to the phospholactyl) group of dehydro-F_420_-0 most likely samples conformations within this pocket where it can be reduced.

Interestingly, the γ-glutamyl ligase CofE from archaea is a single domain enzyme; there is no homology to the C-terminal NTR-fold domain of FbiB. In an analogous situation, Fo synthesis is performed by two single domain enzymes in archaea, CofH and CofG^2^, whereas in bacteria this reaction is catalyzed by a two-domain protein, FbiC (with N-and C-terminal domains homologous to CofH and CofG, respectively). Previous analysis of archaeal genomes revealed that *cofH* and *cofG* are closely associated in genomic context^36^. We therefore investigated the genomic context of archaeal *cofE* genes to investigate whether genes with homology to the C-terminal NTR domain of FbiB were located nearby. From over 1000 archaeal genomes, we only detected 16 open reading frames (ORFs) in the neighboring context of *cofE* that could encode proteins with an NTR fold, although none of these shared substantial (>34% sequence identity) homology and all lacked the key F_420_ binding residues observed in FbiB. There was one interesting exception: the unusual archaea *Lokiarchaeum sp.*^37^ are unique among all sequenced archaea in that they alone encode an FbiB-like γ-glutamyl ligase:NTR fusion protein.

### A recombinant F_420_ biosynthesis operon can be heterologously expressed in *Escherichia coli* to produce mature F_420_

Cofactor F_420_ is only produced by certain bacterial species; the majority of bacteria, including *Escherichia coli,* lack the genes required for F_420_ biosynthesis. Our *in vitro* assay results suggest that 2-PL, and by extension, the hypothesized *L*-lactate kinase CofB, are not required for heterologous production of F_420_ in a non-native organism. To test our hypothesis, we generated a plasmid expressing the *Mycobacterium smegmatis* F_420_ biosynthesis genes *fbiB/C/D* along with the *fbiA* homolog *cofD* from *Methanosarcina mazei*^*19*^, which was substituted as *Ms*-FbiA was found to express poorly in *E. coli*.

We generated two versions of a plasmid-encoded recombinant F_420_ biosynthetic operon, both of which encode codon-optimized genes for expression in *E. coli*: one encodes the native enzymes (pF_420_), and the second encodes C-terminal FLAG-tagged versions of the enzymes to allow their detection in a western blot (pF_420_-FLAG) (Fig. 4a). Plasmids were designed to allow induction of F_420_ biosynthesis in the presence of anhydrotetracycline. A western blot using anti-FLAG antibodies was used to detect whether the proteins were expressed in soluble form in *E. coli*, and confirmed that all were expressed to varying degrees (Fig. S3). We then tested, using HPLC with fluorescence detection, whether F_420_ was heterologously produced in the cell lysate of our recombinant *E. coli* strain expressing this operon. As shown in Fig. 4b, *E. coli* expressing both the FLAG-tagged, and untagged, plasmids produced traces consistent with mature, poly-glutamylated F_420_, although they differed slightly to the *M. smegmatis* produced F_420_ standard in terms of the distribution of poly-γ-glutamate tail lengths.

**Fig. 4.**
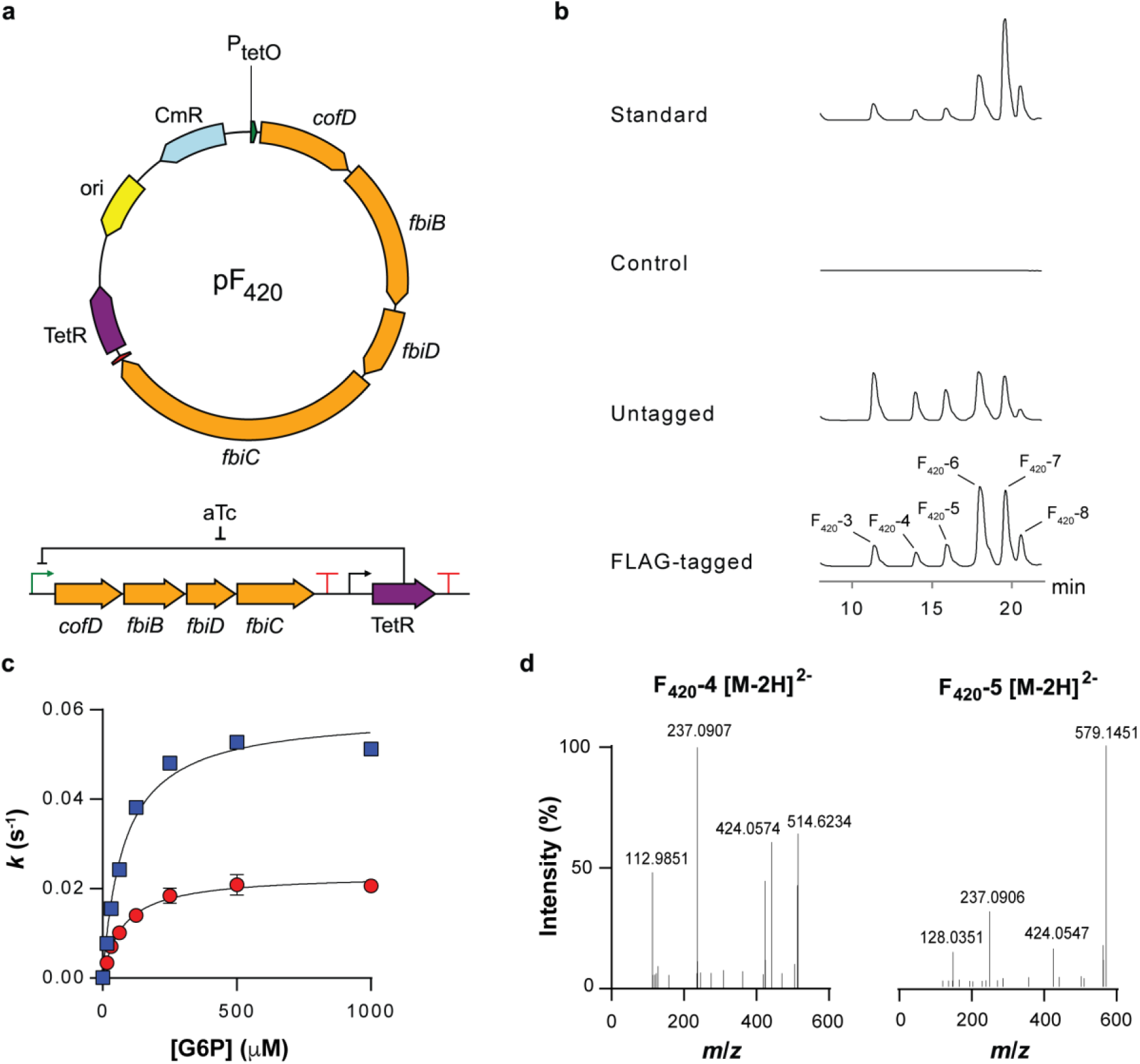
Heterologous expression of F_420_ biosynthesis pathway in *E. coli*. **(a)** Schematic representation of the vector generated for expression of the F_420_ biosynthesis pathway. **(b)** HPLC-FLD traces of *E. coli* lysates containing F_420_ biosynthesis constructs as well as a purified standard from *M. smegmatis.* **(c)** Kinetic studies indicate an identical Michaelis constant for FGD as measured with F_420_ purified from *M. smegmatis* (blue) and *E. coli* (red). **(d)** Fragmentation of F_420_-4 and F_420_-5 extracted from *E. coli* shows a mature F_420_ production.

To confirm that this was indeed mature F_420_, and not dehydro-F_420_ species, we purified the compound from *E. coli* lysate and performed high-resolution MS/MS analysis. Mass fragmentation did indeed show that the compound was reduced, and not dehydro-F_420_ (Fig. 4d). No dehydro-F_420_ species were detected. The yield of purified F_420_ was approximately 27 nmol/L of culture, which is comparable to physiological levels of several F_420_-producing species^38^. UV-Vis and fluorescence spectra of the purified F_420_ matched literature values (Fig. S4)^4,5^. Finally, we confirmed that the purified cofactor was functional by measuring enzyme kinetic parameters with F_420_-dependent glucose-6-phosphate dehydrogenase (FGD) from *M. smegmatis* (Fig. 4c). The apparent Michaelis constant was within error of that observed with FGD and F_420_ produced from *M. smegmatis*, while the *k*cat was approximately half that of the *M. smegmatis* F_420_ (Fig. 4c), which could result from slight differences in the distribution of tail lengths, as previously reported (Table S3)^39^. These results confirm that the recombinant production of F_420_ was achieved with a biosynthetic pathway containing only CofD (equivalent to FbiA)/FbiB/FbiC/FbiD.

## Discussion

It has become widely accepted within the field that one of the essential initial steps in F_420_ biosynthesis involves a hypothetical *L*-lactate kinase that produces 2-PL, which is subsequently incorporated into F_420_ through the activities of CofC and CofD. However, neither bioinformatics nor genetic knockout studies have identified plausible candidate genes for a *L*-lactate kinase^2,18,23^. Furthermore, 2-PL has been shown to inhibit several enzymes involved in central metabolism^28-30^. In terms of pathway flux, this makes 2-PL an unusual starting point for biosynthesis of an abundant metabolite such as F_420_, which can exceed 1 μM in some species^38^. The results presented in this paper unequivocally demonstrate that PEP, rather than 2-PL, is the authentic starting metabolite in bacteria. These results reconcile the previously problematic assumptions that are required to include 2-PL within the biosynthetic pathway and establish a revised pathway (Fig. 5) that is directly linked to central carbon metabolism (*via* PEP) through the glycolysis and gluconeogenesis pathways.

**Fig. 5.**
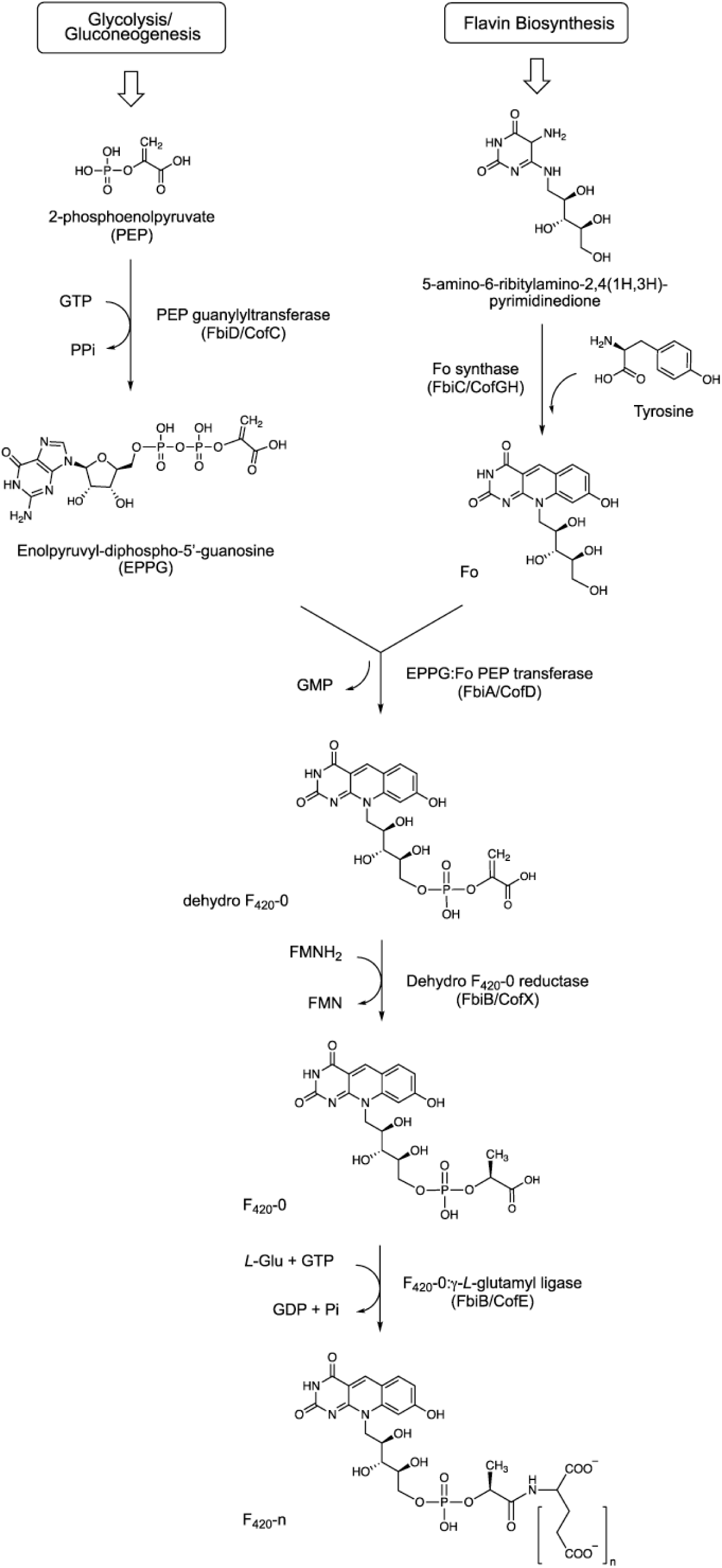
The revised bacterial F_420_ biosynthesis pathway. The revised pathway is a modified scheme showing that PEP acts as the substrate for the FbiD/CofC enzymes to produce EPPG or EPPA (in the case of CofC). The immediate reaction product formed from Fo and EPPG/EPPA is dehydro-F_420_-0, which is reduced to F_420_-0 through the newly-described reductase activity of the C-terminal domain of FbiB in mycobacteria. A separate enzyme in archaea and some bacteria is expected to catalyze this reduction step (CofX). FbiB/CofE subsequently adds a poly-γ-glutamate tail to form F_420_. ‘Fbi’ refers to bacterial proteins, whereas ‘Cof’ represents archaeal ones.

Our observation that CofC accepts PEP (and not 2-PL) *in vitro,* appear to contradict previous studies in which *Mj*-CofC was reported to use 2-PL as substrate^19,33^. However, this discrepancy is most likely due to the supplementation of the coupled CofC/CofD reaction in these studies with pyruvate kinase and 2 mM PEP, a strategy that was used to overcome apparent product inhibition by GMP19,33. Regardless, we cannot explain how a pathway starting from PEP can produce mature (*i.e.* not dehydro-F_420_) F_420_ in archaea given that their equivalent of FbiB (CofE) lacks the C-terminal domain dehydro-F_420_-0 reductase domain seen in FbiB. One possibility is that an unknown dehydro-F_420_-0 reductase exists elsewhere in the genome (remote from CofE). Further studies are required to resolve this step in archaeal F_420_ biosynthesis.

The discovery of dehydro-F_420_-0 as the product of FbiD:FbiA activity in mycobacteria indicated that another enzyme must be required to reduce dehydro-F_420_-0 and produce F_420_-0. This allowed us to define the function of the orphan C-terminal domain of mycobacterial FbiB^20^. The N-terminal domain is homologous with the archaeal γ-glutamyl ligase CofE^33^, but is only capable of catalyzing glutamylation of F_420_-1^20^. The C-terminal domain, which binds both F_420_ and FMN, is essential for extending the poly-γ-glutamate tail of F_420_-0 ^20^, but we could not explain the possible function of the FMN cofactor. Here, we show that FbiB can catalyze both polyglutamylation and reduction of dehydro-F_420_-0, *via* the activities of the N-and C-terminal domains, respectively.

The increasing recognition of the importance of F_420_ in a variety of biotechnological, medical and ecological contexts underlines the need for making the compound more widely accessible to researchers; however our inability to produce F_420_ recombinantly in common laboratory organisms has been a major barrier to wider study. Here, we confirm the results of our *in vitro* experiments by showing that recombinant expression of the four characterized F_420_ biosynthesis genes allows production of F_420_ in *E. coli*. This should now facilitate the use of F_420_ in a variety of processes with recombinant organisms, such as biocatalysis using a bio-orthogonal cofactor, directed evolution of F_420_-dependent enzymes, recombinant production of antibiotics for which F_420_ is a required cofactor, and metabolic engineering.

## Methods

### Bacterial strains and growth conditions

Protein expression utilized either *M. smegmatis* mc^2^4517^40^, *E. coli* BL21(DE3) or LOBSTR-BL21(DE3)^41^ cells. For growth of *M. smegmatis*, media were supplemented to 0.05% (*v*/*v*) Tween80. *M. smegmatis* cells were grown in ZYM-5052^42^ or modified auto-induction media TB 2.0 (2.0% tryptone, 0.5% yeast extract, 0.5% NaCl, 22 mM KH2PO4, 42 mM Na2HPO4, 0.6% glycerol, 0.05% glucose, 0.2% lactose)^43^. For selenomethionine labelling, cells were grown in PASM-5052 media^42^. *E. coli* expressions were conducted in either the above modified auto-induction media, or Terrific Broth (TB) medium modified for auto-induction of protein expression (1.2% tryptone, 2.4% yeast extract, 72 mM K_2_HPO_4_, 17 mM KH2PO_4_, 2 mM MgSO_4_, 0.8% glycerol, 0.015% glucose, 0.5% lactose, 0.375% aspartic acid), grown for 4 h at 37° C followed by overnight incubation at 18° C.

### Protein expression and purification

#### *Mtb*-FbiD

The open reading frame encoding FbiD (Rv2983) was obtained by PCR from *M. tuberculosis* genomic DNA (Table S1). The pYUBDuet-*fbiABD* co-expression construct was then prepared by cloning *fbiD* into pYUBDuet^44^ using *Bam*HI and *Hin*dIII restriction sites, followed by cloning the *fbiAB* operon using *Nde*I and PacI restriction sites. This construct expresses FbiD with an N-terminal His6-tag, whereas the FbiA and FbiB proteins are expressed without any tags.

The pYUBDuet-*fbiABD* vector was transformed into *M. smegmatis* mc^2^4517 strain^40^ for expression. The cells were grown in a fermenter (BioFlo^®^415, New Brunswick Scientific) for 4 days before harvesting. The cells were lysed in 20 mM HEPES, pH 7.5, 150 mM NaCl, 20 mM imidazole, 1 mM β-mercaptoethanol using a cell disruptor (Microfluidizer M-110P) in the presence of Complete protease inhibitor mixture mini EDTA-free tablets (Roche Applied Science). The lysate was centrifuged at 20,000 ×*g* to separate the insoluble material. Recombinant FbiD was separated from other proteins by immobilized metal affinity chromatography (IMAC) on a HisTrap FF 5-ml Ni-NTA column (GE Healthcare), eluted with imidazole, and further purified by Size Exclusion Chromatography (SEC) on a Superdex 75 10/300 column (GE Healthcare) pre-equilibrated in 20 mM HEPES, pH 7.5, 150 mM NaCl, 1 mM β-mercaptoethanol.

#### *Mtb*-FbiA

The open reading frame encoding *M. tuberculosis* FbiA was commercially synthesized and cloned into pRSET-A (Invitrogen). The pYUB28b-*fbiA* construct used for expression in *M. smegmatis* mc^2^4517 was prepared by subcloning *fbiA* into pYUB28b^44^ using an *Nde*I site introduced by overhang PCR utilizing the *Hind*III restriction site present on both vectors amplified with the T7 reverse primer (Table S1). The resulting pYUB28b-*fbiA* construct expresses FbiA with an N-terminal His6-tag. The protein was expressed in *M. smegmatis* mc^2^4517 in ZYM-5052 media auto-induction media^42,45^ in a fermenter (BioFlo^®^415, New Brunswick Scientific) for 4 days. The protein was then purified using Ni-NTA and size-exclusion chromatography steps, as described above, in 20 mM HEPES, pH 7.5, 200 mM NaCl, 5% glycerol, 1 mM β-mercaptoethanol.

#### *Mj*-CofC

The open reading frame encoding *Methanocaldococcus jannaschii* CofC (MJ0887)^25^ was synthesized (GenScript) and cloned into pYUB28b vector^44^ using *Nde*I and *Hin*dIII restriction sites. The protein was expressed in TB auto-induction media and purified using Ni-NTA and size-exclusion chromatography steps, as described above, in 20 mM HEPES, pH 7.5, 200 mM NaCl, 5% glycerol, 1 mM β-mercaptoethanol.

#### *Ec*-Fre

The *E. coli* flavin reductase^35^ was cloned into pProEX-HTb using *Kas*I and *Hind*III restriction sites (Table S1). Protein expression and purification was performed similar to *Mj*-CofC.

#### *Ms*-FbiD

The open reading frame encoding *M. smegmatis* FbiD (MSMEG_2392) was synthesized (Integrated DNA Technologies) and cloned into pETMCSIII by Gibson Assembly as outlined previously^43^. The protein was expressed overnight ay 30 °C in auto-induction media and purified similarly to *Mj*-CofC.

### Construction of synthetic F_420_ biosynthesis operon

Ribosome binding sites were individually optimized for each of the codon-optimized F_420_ biosynthesis genes using the Ribosome Binding Site Calculator server^36,46^. Multiple operon designs were analyzed using the server’s operon calculator and modified to remove unwanted translation products and RNA instability elements while maintaining predicted translation initiation rates for all coding sequences within an order of magnitude. The final design placed the operon under the control of the tetracycline-inducible promoter BBa_R0040 and the artificial terminator BBa_B1006. For subsequent assembly the operon was flanked by BioBrick prefix and suffix sequences. The operon was synthesized by GenScript and cloned into pSB1C3 containing the constitutive tetracycline repressor cassette BBa_K145201 using the standard BioBrick assembly protocol with *Eco*RI and *Xba*I/*Spe*I restriction enzymes^47^.This construct, hereafter referred to as pF_420_, allowed for production of F_420_ to be induced by the addition of anhydrotetracycline. For western blot analysis as second version of the operon with single C-terminal FLAG tags on all four genes was likewise synthesized.

To test for *in vivo* F_420_-depended reductase/oxidase activity the open reading frame of *M. smegmatis* FGD (MSMEG_0777) was codon optimized and commercially synthesized by GenScript and cloned into multiple cloning site 1 of pCOLADuet-1 (Novagen) using *Nco*I and *Hind*III sites. Subsequently MSMEG_2027 was subcloned from pETMCSIII-MSMEG_2027^43^ into multiple cloning site 2 using *Nde*I and *Kpn*I to give pFGD2027.

### *Mtb*-FbiD crystallization and structure determination

*Crystallization*. Apo-FbiD (20 mg/mL in 20 mM HEPES pH 7.5, 150 mM NaCl, 1 mM β-mercaptoethanol) was crystallized using the sitting drop vapor diffusion method in 30% PEG 1500, 3% MPD, 0.2 M MgSO4, 0.1 M sodium acetate pH 5.0. For experimental phasing, selenomethionine-substituted FbiD crystals were grown using protein produced in *M. smegmatis* host cells^45^. Se-SAD anomalous diffraction data sets were collected at the Australian Synchrotron. Data collection statistics are summarized in Table S2. Crystals of *Mtb*-FbiD in complex with PEP were obtained by soaking pre-formed apo crystals in precipitant solutions containing 10 mM PEP for 30 min.

*Structure determination.* All data sets were indexed and processed using XDS^48^, and scaled with AIMLESS^49^ from the CCP4 program suite^50^. The structure was solved using the SAD protocol of Auto-Rickshaw^51^, the EMBL-Hamburg automated crystal structure determination platform. Based on an initial analysis of the data, the maximum resolution for sub-structure determination and initial phase calculation was set to 2.83 Å. All three of the expected heavy atoms were found using the program SHELXD^52^. The initial phases were improved using density modification and phase extension to 2.33 Å resolution using the program RESOLVE^53^. Cycles of automatic model building by ARP/wARP^54^ and phenix.autobuild^55^ resulted in a protein model that was completed manually using COOT^56^. Water molecules were identified by their spherical electron density and appropriate hydrogen bond geometry with the surrounding structure. Following each round of manual model building, the model was refined using REFMAC5^57^, against the data to 1.99 Å resolution. The PDB_redo program^58^ was used in the final stages of refinement. Full refinement statistics are shown in Table S2.

The structure of *Mtb*-FbiD in complex with PEP was solved by molecular replacement using PHASER^59^ with the apo-FbiD structure as a search model. The structure was refined by cycles of manual building using COOT^56^ and refinement using REFMAC5^57^, against the data to 2.18 Å resolution. Full refinement statistics are shown in Table S2.

### HPLC assays

FbiD/CofC-FbiA coupled activity was monitored in a reaction mixture containing 100 mM HEPES pH 7.5, 2 mM GTP, 0.1 mM Fo, 5 mM MgCl_2_, 1 mM 2-PL or PEP, 1 μM FbiD and 5μM FbiA. The reactions were incubated at 37 °C and stopped using 20 mM EDTA at various time points. Separation of F_420_ species was performed on an Agilent HP 1100 HPLC system equipped with photodiode array and fluorescence detectors (Agilent Technologies). Samples were kept at 4°C, and the injection volume was 20 μL. Samples were separated on a Phenomenex Luna C18 column (150 × 3 mm, 5 μm) with a 0.2 μm in-line filter that was maintained at 30 °C. The mobile phase consisted of 100% methanol (A) and 25 mM sodium acetate buffer, pH 6.0 (B), with a gradient elution at a flow rate of 0.5 ml/min and a run time of 30 min. The gradient profile was performed as follows: 0–25 min 95–80% B, 25–26 min 80% B, 26–27 min 95% B, 27–30 min 95% B, and a post-run of 2 min. The wavelengths used for photodiode array were 280 and _420_ nm (bandwidth 20 nm) using a reference of 550 nm (bandwidth 50 nm). The wavelengths used for the fluorescence detector were _420_ nm (excita-tion) and 480 nm (emission).

### LC-MS characterization of dehydro-F_420_ species

Enzymatic reactions were set up as described above. 10 uL aliquots were injected onto a C18 trap cartridge (LC Packings, Amsterdam, Netherlands) for desalting prior to chromatographic separation on a 0.3 × 100 mm 3.5 μm Zorbax 300SB C18 Stablebond column (Agilent Technologies, Santa Clara, CA, USA) using the following gradient at 6 uL/min: 0-3 min 10% B, 24 min 50% B, 26 min 97% B, 29 min 97% B, 30.5 min 10% B, 35 min 10% B, where A was 0.1% formic acid in water and B was 0.1% formic acid in acetonitrile. The column eluate was ionised in the electrospray source of a QSTAR-XL Quadrupole Time-of-Flight mass spectrometer (Applied Biosystems, Foster City, CA, USA). For IDA (Information Dependent Analysis) analyses, a TOF-MS scan from 330-1000 *m/z* was performed, followed by three rounds of MS/MS on the most intense singly or doubly charged precursors in each cycle. For targeted work, defined Product Ion Scans were created to isolate and fragment specific ions of interest with various collision energies (10-60 kV). Both positive and negative modes of ionisation were used as appropriate.

### MS/MS confirmation of reduction of dehydro-F_420_ species

To confirm the reduction of the PEP moiety *in vitro* assays were prepared in 50 mM HEPES pH 7.5, 100 mM KCl, 5 mM MgCl_2_, 2 mM GTP, 0.1 mM Fo, 1 mM PEP, 5 µM *Mtb*-FbiA, 6.5 µM *Msg*-FbiD, 10 mM DTT, 20 µM FMN, 0.2 mM NADH, 0.1 µM *Ec*-Fre, 2 µM *Mtb-*FbiB and 1 mM *L*-glutamate. To minimize futile oxidation of FMNH2 by oxygen the reaction mixture was repeatedly evacuated and purged with nitrogen and maintained under a nitrogen atmosphere. Samples were incubated at ambient temperature for up to 36 hours and stopped by addition of 20 mM EDTA. Samples were desalted using Bond Elut C18 tips (Agilent Technologies) and eluted in 0.1 % formic acid in acetonitrile. Samples were injected on to a Q Exactive Plus (ThermoFischer Scientific) at 100 µl/min with isocratic 50% B where A was 10 mM ammonium acetate pH 6.0 and B was 0.02% ammonia in methanol. Scans from 150-2250 *m/z* were performed and data-dependent MS/MS on targeted metabolites was done in both positive and negative mode.

### Production of F_420_ in *E. coli*

The pF_420_ vector was transformed into *E. coli* BL21(DE3) for expression. Cells were cultivated in M9 minimal media supplemented with chloramphenicol (25 µg/mL) and tyrosine (18 µg/mL) in 1 L shake flasks with 500 ml working volume at 28 °C. At O.D. of 0.60, 200 ng/mL of tetracycline was added to induce the expression of F_420_ biosynthesis pathway. After induction, cells were cultivated for at least 16 h. To test expression of the F_420_ biosynthetic genes, cell pellets were lysed by resuspending to O.D. of 0.8 in buffer containing BugBuster (Novagen). Following centrifugation at 16,000 ×*g* proteins were resolved on a gel (Bolt^TM^ 4-12% Bis-Tris Plus, Invitrogen) for 1 h and visualized using Coomassie Brilliant Blue. For immunoblotting, proteins were transferred to a membrane (iBlot^®^ 2 NC Regular Stacks, Invitrogen) using iBlot Invitrogen (25 V, 6 mins). After staining and de-staining, the membrane was blocked with 3 % skim milk solution and blotted with anti-FLAG antibodies conjugated with HRP (DYKDDDDK Tag Monoclonal Antibody, Thermofisher Scientific).

For detection of F_420_ in *E. coli* lysate by HPLC-FLD, cells were grown overnight in media containing 2.0% tryptone, 0.5% yeast extract, 0.5% NaCl, 22 mM KH_2_PO^4^, 42 mM Na2HPO4, 100 ng/mL anhydrotetracycline, 34 µg/mL chloramphenicol at 30 °C. Cells from 500 µL of culture were pelleted by centrifugation at 16,000 ×*g*. Cells were resuspended in 500 µL of 50 mM Na2HPO4 pH 7.0 and lysed by boiling at 95 °C for 10 minutes. Cell debris was pelleted by centrifugation at 16,000 ×*g* and filtered through a 0.22 µm PVDF filter. Analysis was conducted as described previously^2^. pSB1C3 containing only BBa_K145201 was used as a control.

### Purification and analysis of *E. coli*-produced F_420_

For F_420_ extraction, 1 L of cell culture was centrifuged at 5,000 ×*g* for 15 minutes. The cell pellet was re-suspended in 30 ml of 75% ethanol and boiled at 90 °C in a water bath for 6 min for cell lysis. The cell extract was again centrifuged at 5,000 ×*g* for 15 minutes to remove cell debris and the supernatant was lyophilized. The lyophilized cell extract was re-dissolved in 10 ml milli Q water centrifuged at 5,000 ×*g* for 15 minutes and the supernatant passed through 0.45 µm syringe filter (Millex-HV). The filtered cell extract was purified for F_420_ using a 5 ml HiTrap QFF column (GE Healthcare) as previously described^44^. The purified F_420_ solution was further desalted by passing it through C18 extract column (6 ml, HyperSep C18 Cartridges, ThermoFischer Scientific). The C18 extract column was first equilibrated by passing through 10 ml of 100% methanol and 10 ml of milli Q. Afterwards the F_420_ solution was passed through the C18 extract column, F_420_ was eluted in 2 ml fraction of 20% methanol.The purified F_420_ solution was further concentrated in GeneVac RapidVap and re-dissolved in 500 µl of milliQ for further analysis and assays.

UV and fluorescence spectra were collected on a Varian Cary 60 and a Varian Cary Eclipse, respectively, in a 10 mm QS quartz cell (Hellma Analytics). Samples were buffered to pH 7.5 with 50 mM HEPES and scanned from 250-600 nm. For fluorescence, the excitation wavelength was _420_ nm and the emission scanned from 435 to 600 nm.

Activity assays with *M. smegmatis* and *E. coli*-derived F_420_ were conducted with *M. smegmatis* FGD1 expressed and purified as described previously^15^. Assays were performed in 50 mM Tris pH 7.5, 300 mM NaCl, 50 nM FGD1, 5 µM F_420_ and 0-1000 µM glucose-6-phosphate using a SpectraMAX e2 plate reader. Activity was measured by following loss of F_420_ fluorescence at 470 nm. Apparent *k*cat and *K*m values were calculated using GraphPad Prism 7.04 (GraphPad Software, La Jolla California).

### 2-PL synthesis

In the absence of a commercial source, 2-PL was chemically synthesized by a slight modification of the method of Kirsch^60^. Briefly, benzyl lactate was condensed with chlorodiphenyl phosphate in pyridine, with cooling, to give benzyldiphenylphosphoryl lactate. Hydrogenolysis of this material in 70% aqueous tetrahydrofuran over 10% Pd-C gave phospholactic acid as a colorless, viscous oil, which was characterized by proton, carbon and phosphorus NMR spectroscopy, and by mass spectrometry. ^1^H NMR (DMSO-d6) δ 11.68 (br, 3H), 4.53 (m, 1H), 1.36 (d, J=6.8 Hz, 3H). ^13^C NMR (DMSO-d6) δ 172.50 (d, JP-C=0.05 Hz), 69.56 (d, JP-C=0.04 Hz), 19.27 (d, JP-C=0.04 Hz). ^31^P NMR (DMSO-d6) δ −1.64. APCI-MS found: [M+H]^+^=171.1, [M-H]^-^=169.1.

### Fo purification

Fo was purified from *M. smegmatis* culture medium over-expressing *Mtb*-FbiC as described previously^44^.

### Genomic context analysis

A non-redundant CofE dataset of 4,813 sequences was collected using the Pfam identifier PF01996 and the InterPro classifications IPR008225, IPR002847, and IPR023659. Archaeal CofE sequences were extracted from this dataset, resulting in a set of 1,060 sequences that included representatives across twelve phyla: Crenarchaeota, Euryarchaeota, Thaumarchaeota, *Candidatus* Bathyarchaeota, *Candidatus* Diapherotrites, *Candidatus* Heimdallarchaeota, *Candidatus* Korarchaeota, *Candidatus* Lokiarchaeota, *Candidatus* Marsarchaeota, *Candidatus* Micrarchaeota, *Candidatus* Odinarchaeota, and *Candidatus* Thorarchaeota. The genomic context (10 upstream and 10 downstream genes) of each archaeal *CofE* was analyzed for the presence of a neighboring PF00881 domain with homology to the C-terminal domain of FbiB using the Enzyme Function Initiative-Genome Neighborhood Tool^61^.

## Acknowledgment

We thank Assoc. Prof. Chris Squire, Dr Carol Hartley and Dr Andrew Warden, and Dr Matthew Taylor for helpful discussions, Dr Adam Carroll and Dr Thy Truong for technical assistance. This research is supported by a Sir Charles Hercus Fellowship (GB), an NHMRC New Investigator Grant (CG; 1142699), an ARC DECRA Fellowship (CG; DE170100310), the Health Research Council of New Zealand (ENB), an AGRTP Scholarship (JA), the Australian Research Council, and the National Health and Medical Research Council (CJ). Crystal data collection was undertaken on the MX1 beamline at the Australian Synchrotron (Victoria, Australia). Access to the Australian Synchrotron was supported by the New Zealand Synchrotron Group Ltd.

## Author contribution

JA performed experiments, analysed results and cowrote the manuscript; ENMJ, MH, JC, SMS, SS, BP, MM performed experiments and analyzed results; BN conceived project, performed experiments and analyzed results; CG, ENB conceived project, designed experiments, analyzed results and cowrote the manuscript; CS designed experiments, analyzed results and cowrote the manuscript; GB, CJJ conceived project, designed experiments, performed experiments, analyzed results and cowrote the manuscript. All authors provided feedback on the manuscript.

---

**Fig. S1.**
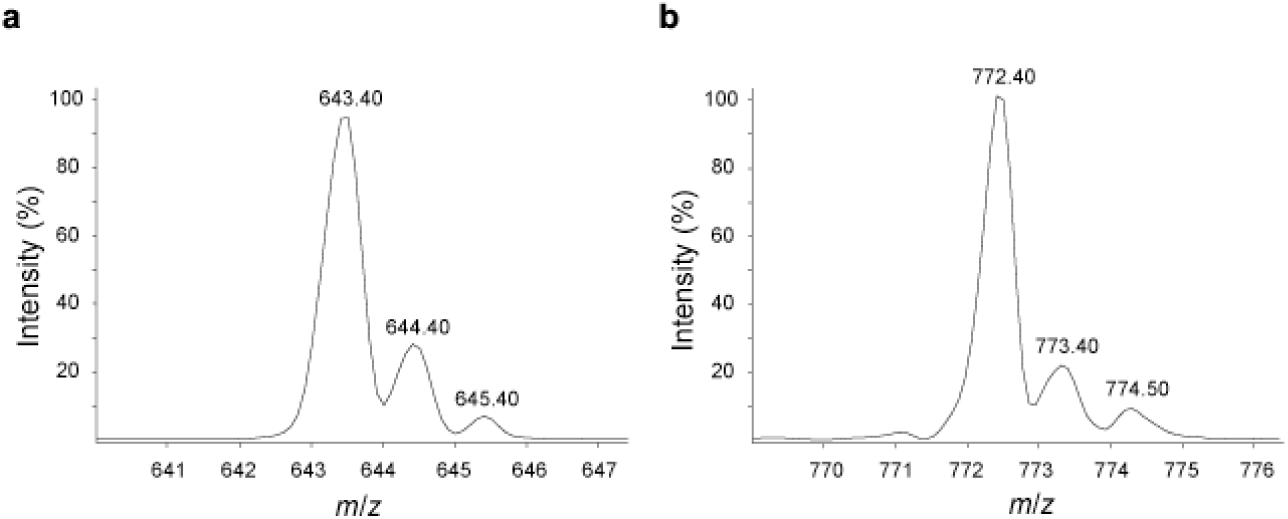
Addition of *L*-glutamate residues to dehydro-F_420_-0. Dehydro-F_420_-1 ([M+H]^+^, monoisotopic *m*/*z* of 643.40) and dehydro-F_420_-2 ([M+H]^+^, monoisotopic *m*/*z* of 772.40) are formed upon the addition of one and two *L*-glutamate residues, respectively, to dehydro-F_420_-0.

**Fig. S2.**
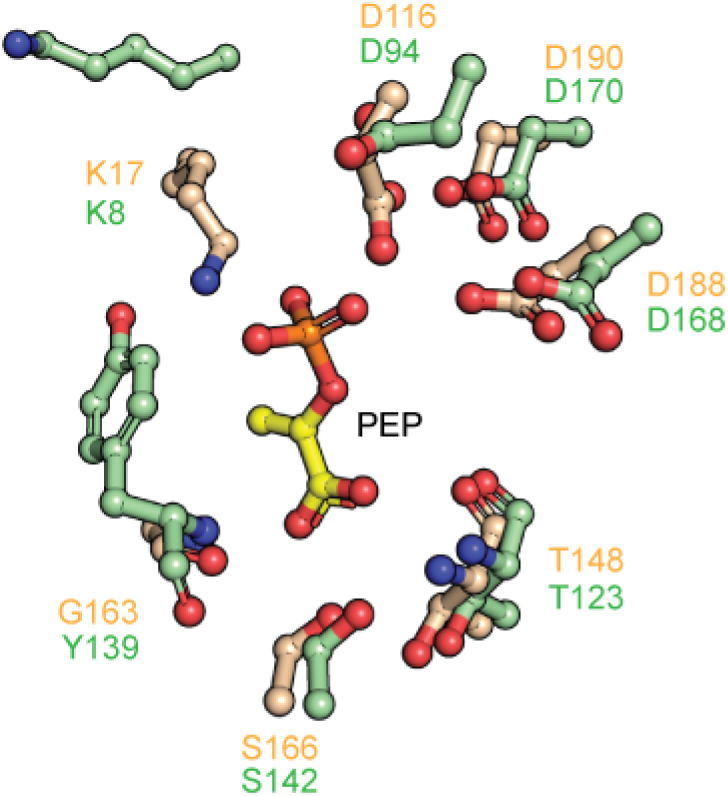
Conservation of PEP binding residues between *Mtb*-FbiD and *Mj*-CofC. Superposition of the *Mtb*-FbiD (wheat) on to that of *Mj*-CofC (green) indicates conservation of the residues in the PEP binding site of both proteins. The only change in the binding site, Gly>Tyr in *Mj*-CofC, is not likely to affect binding considering that the hydrogen bond interaction takes place between the backbone nitrogen atom and the oxygen atom of PEP carboxylate group. Protein side chains are shown in ball-and-stick model and labelled with the corresponding color.

**Figure S3.**
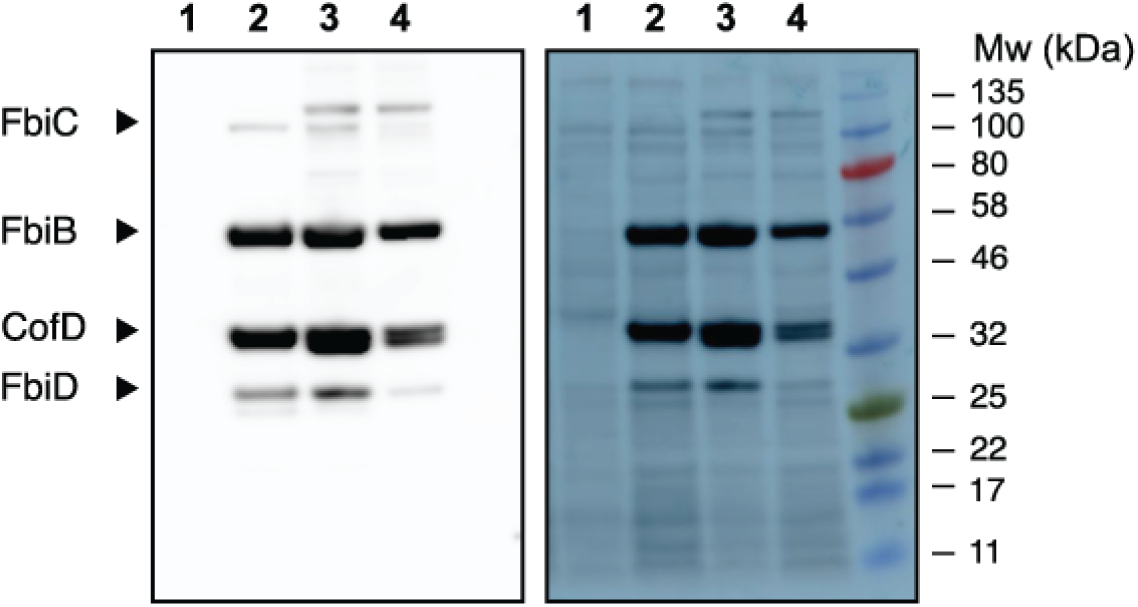
Expression of F_420_ biosynthetic genes in *E. coli*. Whole cell lysates were re-solved on SDS-PAGE, then immunoblotted with anti-FLAG antibodies to detect expression of F_420_. 1) Vector-only control 28 °C; 2) pF_420_-FLAG expressed at 37 °C; 3) pF_420_-FLAG expressed at 28 °C; pF_420_-FLAG expressed at 18 °C.

**Figure S4.**
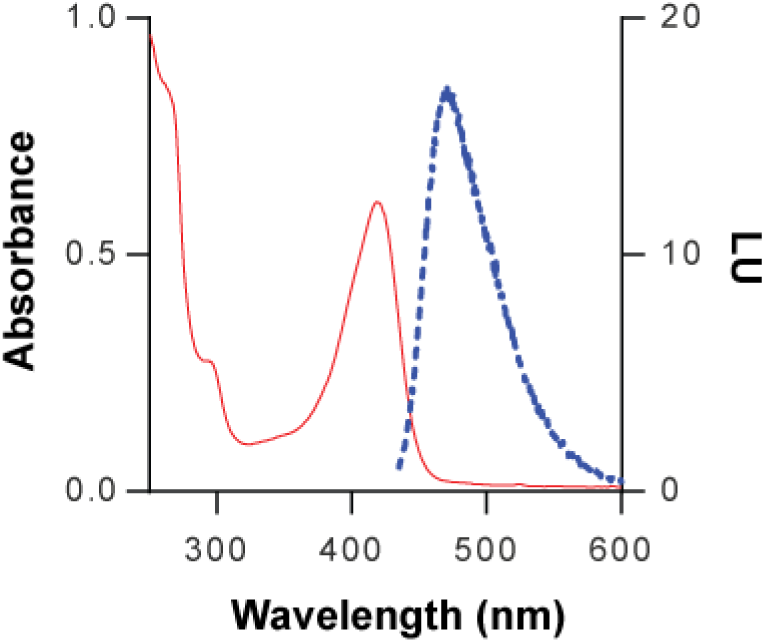
Spectrophotometric identification of *E. coli*-produced F_420_. UV-Vis scan (red) and fluorescence emission (blue) spectra indicate the characteristic features of F_420_; maximum absorption at _420_ nm and maximum fluorescence emission at 480 nm.

**Table S1.**
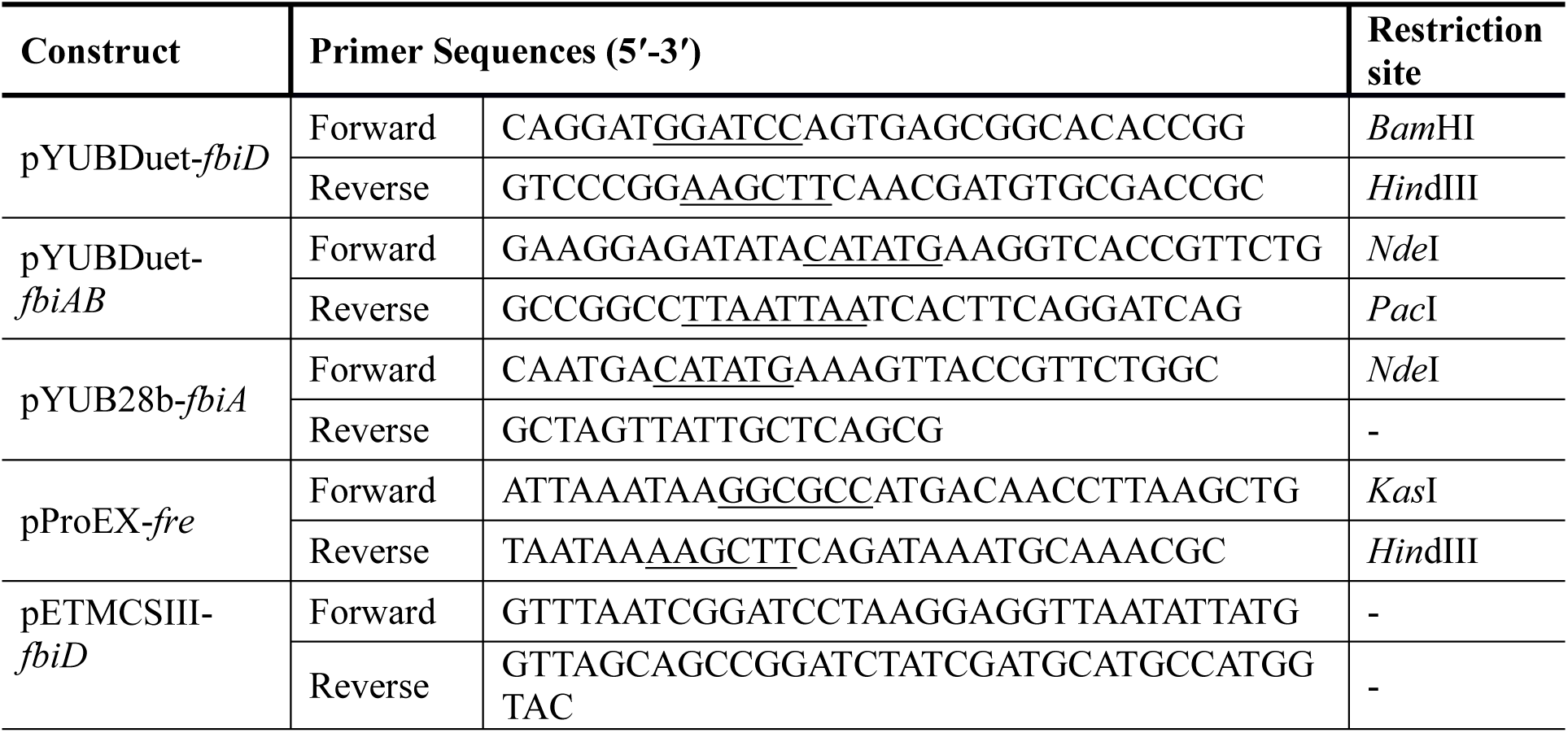
Primers used in the amplification of the constructs used in this study.

**Table S2.**
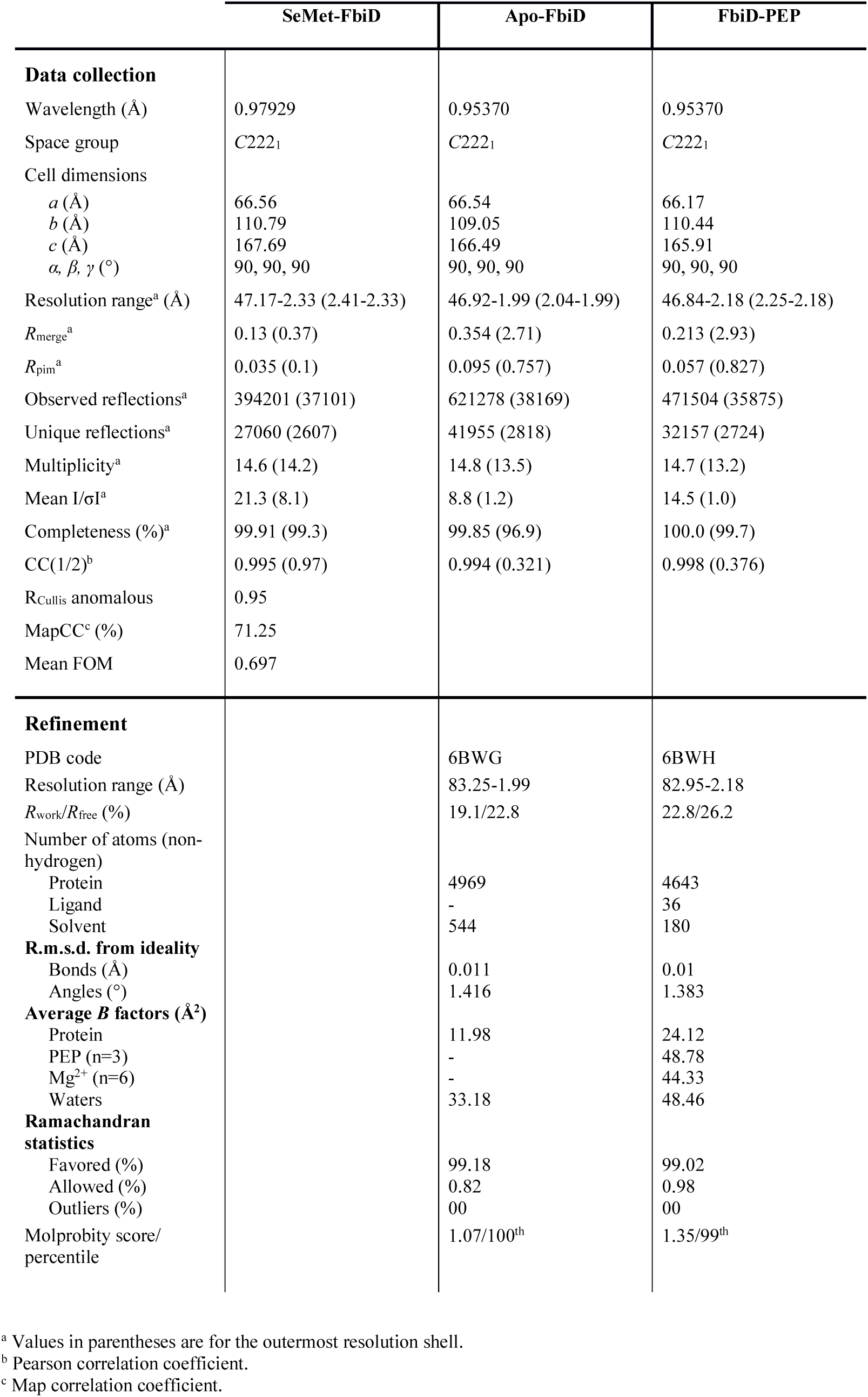
Data collection, processing and refinement statistics.

**Table S3.** Kinetic parameters of FGD with F_420_ purified from *M. smegmatis* and *E. coli*.

| | $k_{\text{cat (app)}} \text{ (s}^{-1}\text{)}$ | $K_{\text{m (app)}} \text{ (}\mu\text{M)}$ |
| --- | --- | --- |
| *M. smegmatis* | $1.19 \pm 0.05$ | $79 \pm 11$ |
| *E. coli* | $0.46 \pm 0.02$ | $77 \pm 7$ |

